# *LsBADH1* is responsible for sweet fragrance in lettuce (*Lactuca sativa* L.) through 2-acetyl-1-pyrroline biosynthesis

**DOI:** 10.64898/2026.06.13.731611

**Authors:** Kousuke Seki, Kenji Matsui, Miya Yanagidate, Keiji Nishida, Ryohei Koyama, Yuichi Uno

**Affiliations:** Nagano Vegetable and Ornamental Crops Experiment Station, Tokoo, Souga, Shiojiri, Nagano 399-6461, Japan; Graduate School of Sciences and Technology for Innovation, Yamaguchi University, Yamaguchi, 753-8515, Japan; Plant Science Division, Department of Bioresource Science, Graduate School of Agricultural Science, Kobe University, Rokkodai 1-1, Nada, Kobe, Hyogo, 657-8501, Japan; Engineering Biology Research Center, Kobe University, 7-1-49, Minatojima Minami Machi, Chuo-ku, Kobe 650-0047, Japan

**Keywords:** 2-acetyl-1-pyrroline, betaine aldehyde dehydrogenase, fragrance, volatile compounds, lettuce, stem type

## Abstract

**Highlight:** The sweet fragrance of lettuce was attributed, for the first time, to the synthesis of 2-acetyl-1-pyrroline caused by a deficiency in the betaine aldehyde dehydrogenase gene.

Fragrance is among the most valuable traits of high-quality crops and influences consumer preferences. Although 2-acetyl-1-pyrroline (2AP) is a key component of fragrant cultivars in several crops, its genetic mechanism in lettuce (*Lactuca sativa* L.) remains poorly understood. The betaine aldehyde dehydrogenase (*BADH*) gene has been identified as causative for 2AP-derived fragrance in rice and soybean cultivars. Hence, we conducted a linkage analysis using an F_2_ population derived from a cross between ‘Kukichisya’ (fragrant) and ‘Rennet’ (non-fragrant) for three candidate genes of *BADH* orthologs in the lettuce genome. Analysis linked *LOC111877932* located in LG4 to the fragrance trait, and it was designated *LsBADH1*. Comparison among ‘Kukichisya’, ‘Salinas’, and candidate *BADH* of sunflower (*Helianthus annuus* L.) revealed three non-synonymous single-nucleotide polymorphisms (nsSNPs) in exons 1, 2, and 9, and suggested that nsSNP in exon 9 was strongly correlated with fragrance in ‘Kukichisya.’ A premature stop codon introduced in exon 5 of *LsBADH1* using Target-AID base-editing technology resulted in truncated BADH1 and higher 2AP levels. Our results indicated that *LsBADH1* is responsible for the 2AP-derived fragrance. Our findings can be applied to select cultivars based on a novel concept for the cooking process, providing a transformative platform to breed fragrant lettuce as a high-value-added product.

## Introduction

Several horticultural types of lettuce are cultivated worldwide and are generally categorized based on head and leaf shape and structure and stem shape. The five main types include crisphead, leaf (green and red), romaine, butterhead, and stem. While most types are consumed as fresh vegetables in salads (Ryder, 1999), the stem type is typically used in Chinese cuisine after being peeled or dried. Recently, we developed the pioneer fragrant cultivar ‘Hisui no Kaori’, which has a strong sweet fragrance, by crossbreeding the fragrant stem-type ‘Kukichisya’ with the non-fragrant‘Rennet’ (Seki *et al*., 2024). The characteristic traits of this fragrant type are a strong sweet fragrance, edible leaves and stems, and fresh green color that persists after heat cooking. The diversity of lettuce leaves with regard to textures such as crispness is widely recognized, but fragrance is a surprising trait among leafy vegetables, and could expand the culinary potential of lettuce. The fragrant type was demonstrated to contain the volatile compound 2-acetyl-1-pyrroline (2AP), which has been reported to accumulate in the high-quality fragrant cultivars of soybean (*Glycine max* (L.) Merr.) and rice (*Oryza sativa* L.) (Paule and Powers, 1989; Wu *et al*., 2009; Arikit *et al*., 2011). Fragrance is considered one of the most important quality traits of vegetables because it is a key factor in determining market prices (Somta *et al*., 2019). Previous studies have indicated that orthologs to betaine aldehyde dehydrogenase (*BADH*) genes are associated with the fragrance arising from 2AP accumulation in soybeans, rice, sorghum, and cucumbers (Chen *et al*., 2008; Kovach *et al*., 2009; Arikit *et al*., 2011; Juwattanasomran *et al*., 2011; Yundaeng *et al*., 2013, 2015). Therefore, *BADH* is hypothesized to be the candidate gene responsible for fragrance in lettuce. To test this hypothesis, we investigated the association between the candidate gene orthologous to *BADH* and fragrance using the F_2_ population. In addition, we analyzed the whole-genome sequence of the fragrant ‘Kukichisya’ to identify the gene responsible for fragrance, and conducted genome editing to verify whether the candidate gene is responsible for 2AP-derived fragrance in lettuce.

## Materials and Methods

### Plant materials

‘Kukichisya’ is a fragrant cultivar in our genetic resources (Seki *et al*., 2024), while ‘Celtuce’ and ‘Rennet’ are non-fragrant cultivars. For identifying fragrance-associated compounds using gas chromatography-mass spectrometry (GC-MS), ‘Kukichisya’ and ‘Celtuce’ were cultivated from June 2012 to September 2012 at Nagano Vegetable and Ornamental Crops Experiment Station (Shiojiri city, Nagano prefecture, Japan; 36° 10’ N, 137° 93’ E). For linkage analysis, 188 F_2_ individuals were obtained by a cross between ‘Kukichisya’ and ‘Rennet.’ For genome editing, the cultivar ‘Oilseed’ was used due to its short life cycle and capacity for rapid generation advancement.

### Linkage analysis

One hundred and eighty-eight individuals of the F_2_ population were derived from a cross between ‘Kukichisya’ (fragrant) and ‘Rennet’ (non-fragrant). The F_2_ individuals and their parents were grown from September 2013 to March 2014 in a greenhouse at Nagano Vegetable and Ornamental Crops Experiment Station for DNA extraction and fragrance evaluation by organoleptic testing using fresh leaves. Genomic DNA was extracted from young leaves using the NucleoSpin Plant II Extract Kit (Machery–Nagel, Duren, Germany).

### Development of PCR-based markers

Two genes, *GmBADH1* (Glyma06g19820) and *GmBADH2* (Glyma05g01770), coding *BADH* in soybean (Juwattanasomran *et al*., 2011) were searched against the lettuce reference genome sequence [version8 from crisphead cultivar ‘Salinas’ (https://genomevolution.org/coge/GenomeInfo.pl?gid=28333)] using GBrowse at the Lettuce Genome Resource (LGR: https://lgr.genomecenter.ucdavis.edu/Home.php) to identify candidate *BADH* orthologous genes (Reyes-Chin-Wo *et al*., 2017). Simple sequence repeats (SSRs) were searched for in the predicted genes and the DNA sequences surrounding the genes using the SSRIT program (http://www.gramene.org/microsat/ssr.html). Primers for amplifying the markers were designed using Primer3 (http://bioinfo.ut.ee/primer3-0.4.0/), and their IDs (names) were defined as (Linkage group)_(genome version)_(genome position). For further genetic mapping to narrow down the gene position, polymorphisms near the genomic region linked to the fragrance trait were identified as marker sites using the IGV software (Robinson *et al*., 2011). PCR was performed using 0.5□μL DNA template, 0.4□μL of each primer (50□μM), 2□μL dNTPs (2□mM), 5□μL 2× PCR Buffer, 0.2□μL KOD FX (1□U μL^−1^, TOYOBO, Japan), and distilled water (dH_2_O) to a final volume of 10□μL. PCR conditions were as follows: 94□°C for 5□min, 30 cycles of 94□°C for 30□s and 61□°C for 30□s, followed by one cycle at 72□°C for 4□min. After amplification, 9 μL of the PCR products were electrophoresed on 2% agarose gel (Takara-bio, Japan) at 100 V. Genetic relationships among the markers were calculated using the AntMap program (Iwata and Ninomiya, 2006), and the Kosambi mapping function was used to calculate the distances between the marker loci in cM. The linkage map was graphically visualized using MapChart (Voorrips, 2002).

### Phylogenetic analysis

BADH protein sequences were used for phylogenetic analysis. Multiple sequence alignments of the full-length protein sequences were performed using ClustalW (https://embnet.vital-it.ch/software/ClustalW.html). A phylogenetic tree was generated using the MEGA X program using the neighbor-joining method with default parameters and 1,000 bootstrap replications (Kumar et al. 2018).

### Re-sequencing analysis

Genomic DNA was extracted from young leaves of ‘Kukichisya’ using NucleoSpin Plant II (Machery–Nagel, Duren, Germany), and was used to construct paired-end sequencing libraries (150 bp ×2) and subjected to whole-genome sequencing using the HiSeq X platform. Re-sequencing analyses were performed as described previously (Seki *et al*., 2020). Raw sequence data (fastq) for the present analysis are available in the DDBJ Sequence Read Archive under accession number DRA012781.

### Genome editing

A 20-nt gRNA specific for the target gene *LsBADH1* was designed at the 5^th^ exon to introduce the stop codon TAA by changing the original tryptophan codon (TGG) using Target-AID base editing (Fig. S1) (Nishida *et al*., 2016). An oligo DNA pair containing the target sequence flanked by the *Bbs*I ligation overhangs was annealed and ligated into the vector, 1480_MluI-1433_pUC19_AtU6oligo (Shimatani *et al*., 2017). The gRNA expression cassette was excised using I-SceI and subcloned into the dicot-optimized Target-AID T-DNA vector 4427_9_pZD_AtU6_HolgerCas9 (D10A) AtPmCDA1_UGI_NPTII (Shimatani *et al*., 2017). All vectors were constructed by standard cloning and verified by sequencing. Lettuce ‘Oil seed’ was transformed using *Agrobacterium* and the transformed plants were selected using kanamycin; genome integration of T-DNA was confirmed by using PCR amplifying *Cas9* and *NPTII* regions. To analyze the outcomes of genome editing, genomic DNA was extracted from individual plants and the *LsBADH1* locus containing the target sequence was amplified using PCR. PCR fragments were cloned into the standard sequencing vector pSKII and analyzed using Sanger sequencing. Eight plasmid clones from each strain were analyzed to determine their homogeneity.

### Evaluation of fragrance-related compounds

Leaf of ‘Kukichisya’ and ‘Celtuce’ was evaluated for fragrance-related compounds using GC-MS. The aerial plant organs (2.5 g fresh weight) were cut into pieces, and sealed in a glass vial (22 mL; Perkin Elmer, Waltham, MA, USA), and frozen at −80 °C for at least 24 h. The glass vial with the plant material was immersed into a water bath set at 25 °C for 10 min. After thawing the plant materials, an SPME fiber (50/30 µm DVB/Carboxen/PDMS, Supelco, Bellefonte, PA, USA) was exposed to the headspace of the vial for 30 min at 25 °C. The fiber was inserted into the insertion port of a GC-MS system (QP-5050, Shimadzu, Kyoto, Japan) equipped with a 0.25 µm × 30 m Stabiliwax column (Restek, Bellefonte, PA, USA). The column temperature was programmed as 40 °C for 1 min, followed by an increas of 15 °C min^−1^ to 180 °C, at which it was held for 1 min. The carrier gas (He) was delivered at a flow rate of 1 mL min^−1^. An SPME Sleeve (Supelco) was used as the glass insert. Splitless injection with a sampling time of 1 min was used. The fiber was held in the injection port for 10 min to completely remove any compounds from the matrix. The temperatures of the injector and interface were 200 °C and 230 °C, respectively. The mass detector was operated in the electron impact mode with an ionization energy of 70 eV. To identify and confirm 2AP, the leaves of *Pandanus amaryllifolius* were analyzed under the same GC-MS conditions (Wakte *et al*., 2010), and the retention time and the MS profiles were compared with those obtained with the ‘Kukichisya.’ Student’s *t*-test was used to determine significant differences between the two cultivars.

### Headspace-solid phase microextraction combined with GC-MS analysis

Volatile compounds were obtained by using headspace solid-phase microextraction (HS-SPME). The GC-MS/MS system (TQ-8040NX) was equipped with an AOC-6000 autosampler (Shimadzu, Kyoto, Japan). For each analysis, 20 g of cut lettuce petiole was placed in a 20 mL headspace vial. Subsequently, the samples underwent equilibration at 90 °C for 15 min in AOC-6000 system. Extraction was conducted at 90 °C using a divinylbenzene/polydimethylsiloxane (DVB/PDMS), a smart SPME Arrow fiber (120 μm film thickness, Shimadzu), inserted into the headspace of each vial for 10 min. Volatile compoundson the SPME Arrow fiber were desorbed for 2 min at 200 °C in the GC splitless injector. The volatile compounds were isolated on a DB-5 capillary column (30 m × 0.25 mm × 0.25 μm, Agilent Technologies, Santa Clara, CA, USA). Helium served as the carrier gas, and the flow velocity was set to 39.0 cm s^−1^. The initial temperature program remained at 40 °C for 5 min. The column temperature was raised from 40 °C to 80 °C at a rate of 5 °C min^−1^, and then to 200 °C at a rate of 15 °C min^−1^, and then maintained at 200 °C for 5 min. The standard ionization energy for routine GC–MS methods in the electron ionization (EI) mode is 70 eV. The mass scan range was 35–350 *m*/*z*. The ion source and ion source temperatures were set to 200 °C and 240 °C, respectively. 2AP was purchased from Toronto Research Chemicals and used as the standard compound. The other compounds were tentatively identified based on the NIST database and a comparison with the RI found in the literature.

## Results

### Inheritance of fragrance trait derived from ‘Kukichisya’

Fragrant genotypes of the F_2_ individuals from a cross between ‘Kukichisya’ (fragrant) and ‘Rennet’ (non-fragrant) were determined by an organoleptic test using fresh leaves grown in the greenhouse. Fragrance derived from ‘Kukichisya’ was suggested to be controlled by a single recessive gene, according to segregation of putative genotype of causative gene showing 1:3 ratio in the F_2_ population (Table 1).

**Table 1.**
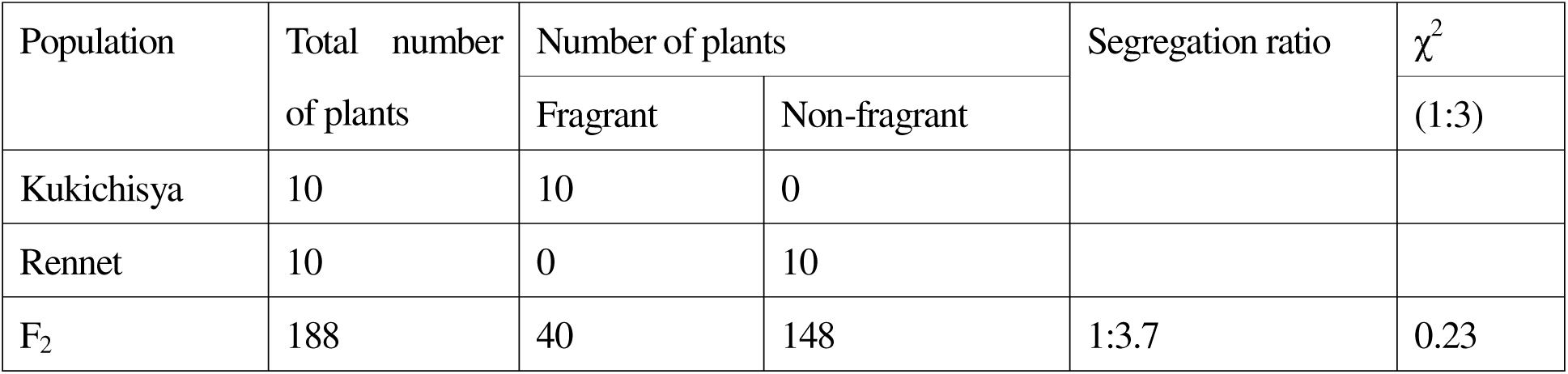
Segregation of the fragrance in F_2_ population derived from ‘Kukichisha’ and ‘Rennet’.

### Linkage analysis with SSR and InDel markers

Using GBrowse at LGR, three gene models were identified as homologous to *GmBADH1* but not *GmBADH2*. Phylogenetic analysis of the three *BADH1* genes in soybean, rice, and sorghum showed that the three gene models of lettuce were categorized into the BADH1 group (Fig. 1) and were considered candidates for the causative gene. The details of three gene models were described below and were based on the latest version 15 of the lettuce reference genome sequence. *LOC111877932* (alias gene ID:111877932) was located on LG 4 from 125,128,259 to 125,132,307 bp (-strand), with a full genomic sequence length of 3,236 bp. The predicted protein of *LOC111877932* contained 504 amino acids translated from 15 exons with a total coding sequence (CDS) of 1,515 bp. *LOC111897919* (alias gene ID:111897919) was on LG5 positioned from 244,613,666 to 244,617,548 bp (-strand), with a full genomic sequence length of 3,578 bp. The predicted protein of *LOC111897919* was 495 amino acids in length, being translated from nine exons with a total CDS length of 1,488 bp. *ALDH10A9* (alias gene ID:111881493) was on LG 5 positioned from 40,430,325 to 40,432,420 bp (+ strand), with a full genomic sequence length of 1,293 bp. The predicted protein of, *ALDH10A9*was 173 amino acids in length, being translated from six exons with a total CDS length of 522 bp. Three SSR markers linked to three gene models, *LG4_v15_125.393Mbp*, *LG5_v15_40.221Mbp*, and *LG5_v15_244.540Mbp*, were developed using reference genome sequences v15 (Table 2).

**Fig. 1.**
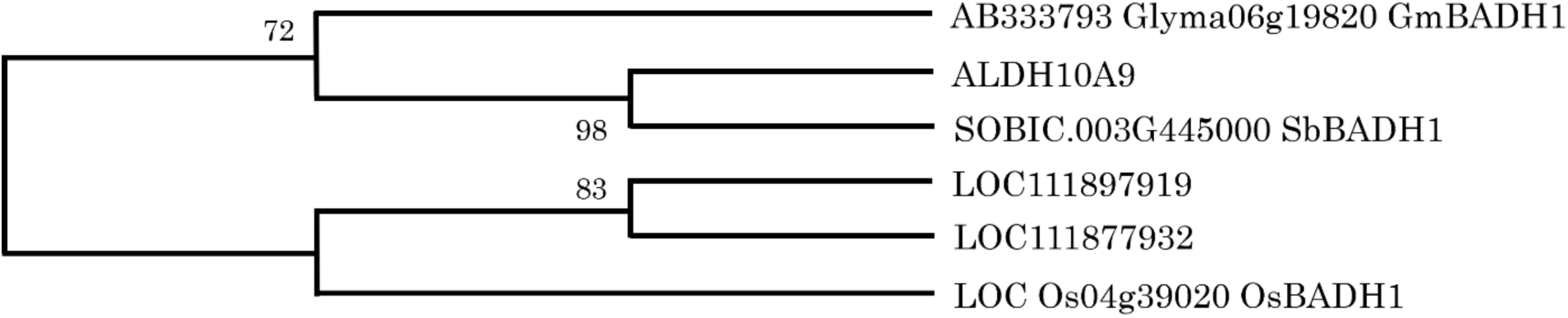
A neighbor-joining phylogenetic tree of three gene models of lettuce and *BADH1* homologs in some plants. Bootstrap values are the percentage of 1,000 replicates.

**Table 2.**
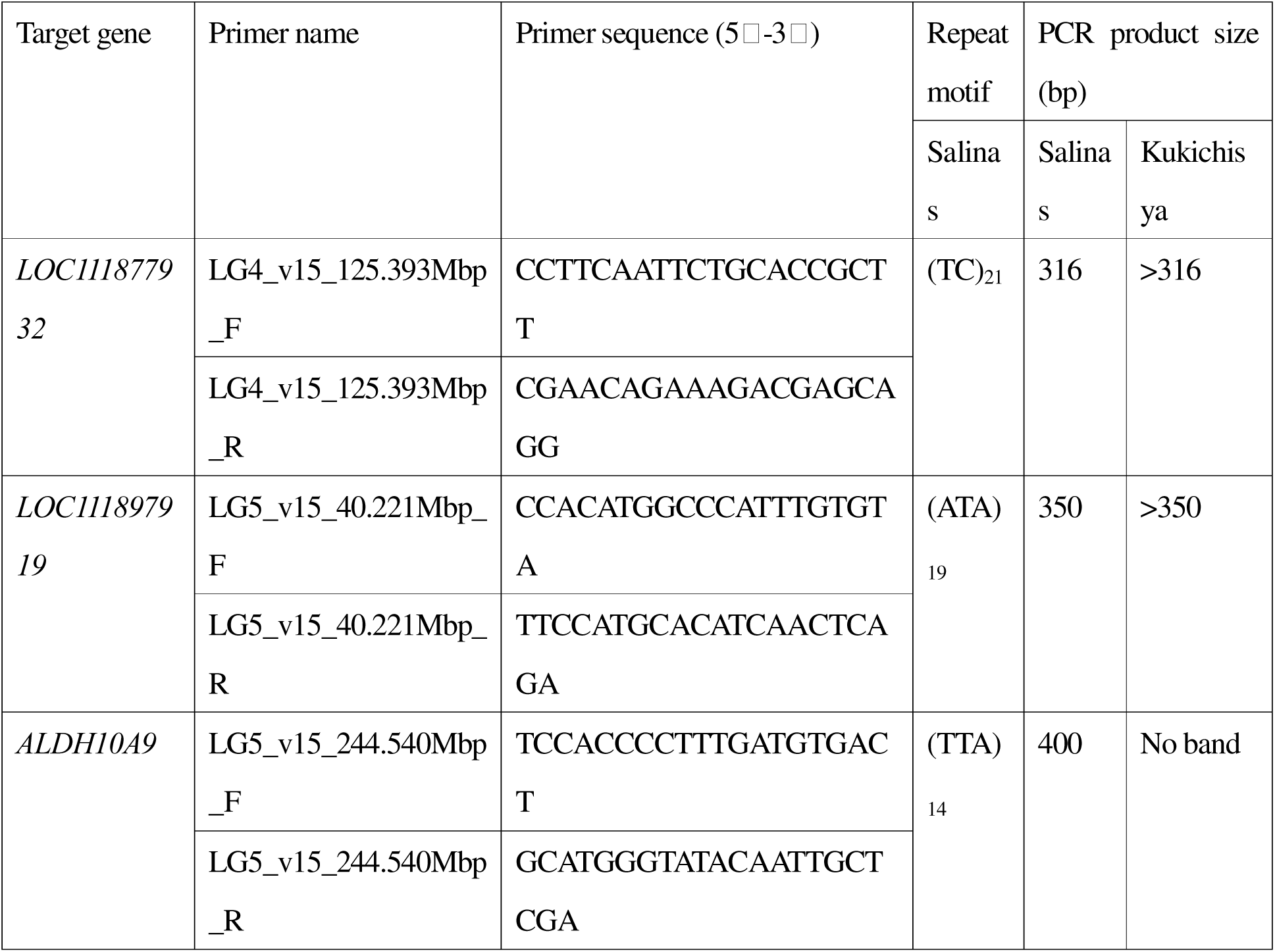
Simple sequence repeat markers designed for detecting genome polymorphism in the vicinity of three *BADH* candidate genes.

Linkage analysis using 188 F_2_ individuals revealed that the SSR primer *LG4_v15_125.393Mbp* was mapped at a distance of 0.3 cM from the causative gene of fragrance, and *LG5_v15_40.221Mbp* and *LG5_v15_244.540Mbp* were unlinked (Fig. 2). Further linkage analysis to narrow down the position of the causative gene in the 2AP locus using several PCR-based markers revealed that the causative gene was located between *LG4_v15_124.067Mbp* and *LG4_v15_125.392Mbp*, and *LG4_v15_124.375Mbp* and *LG4_v15_125.130Mbp* showed complete co-segregation with the fragrance owing to 2AP (Fig. 2, Table 3). Because *LOC111877932* was also located at the same position and was considered a biologically plausible candidate gene for fragrance caused by 2AP, it was designated *LsBADH1*.

**Fig. 2.**
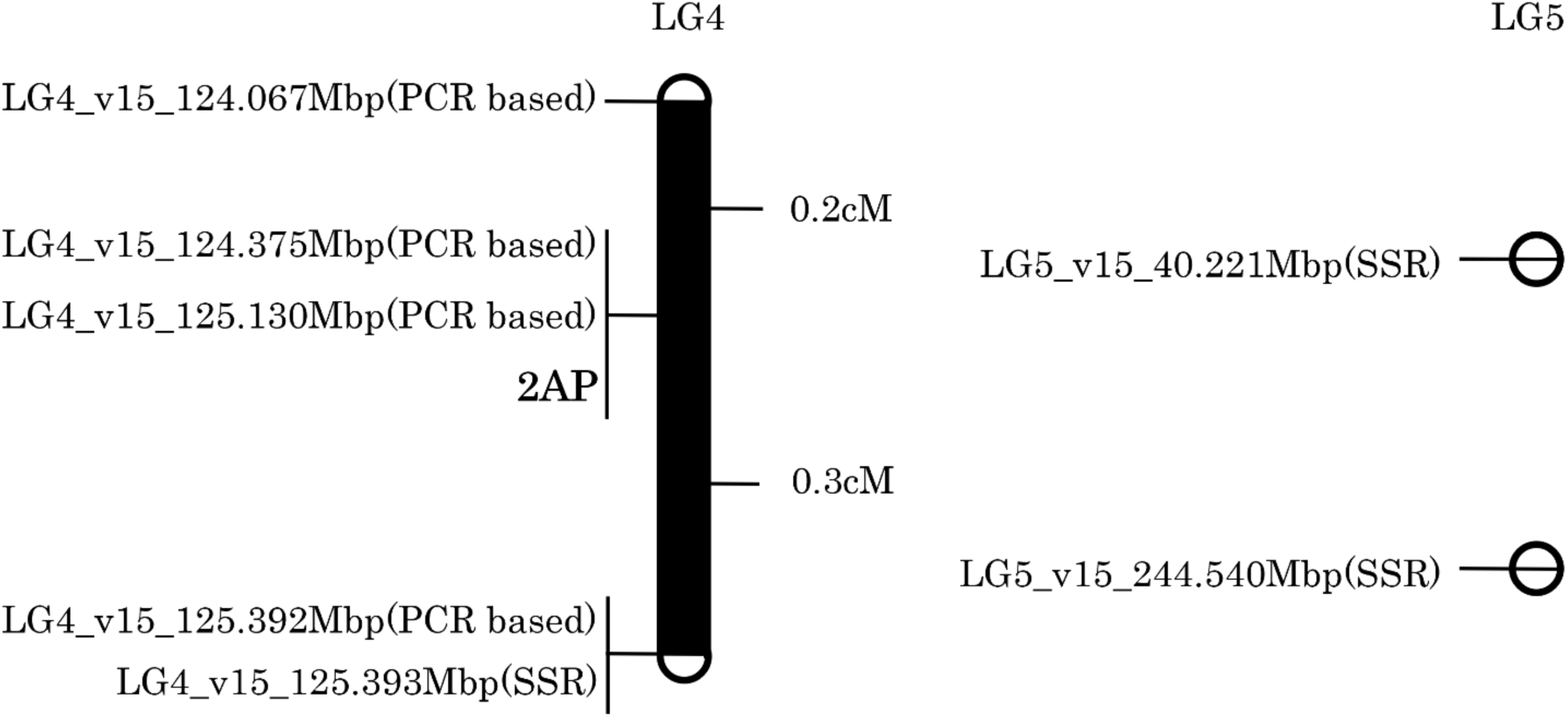
Fine mapping of the 2-acetyl-1-pyrroline (2AP) locus on LG4. Genetic distances (cM) are shown between the markers. “(SSR)” and “(PCR based)” in the marker name indicate simple sequence repeat (SSR) markers and PCR-based markers, respectively. “2AP” in the map indicates the position of the causal gene for fragrance owing to 2AP.

**Table 3.**
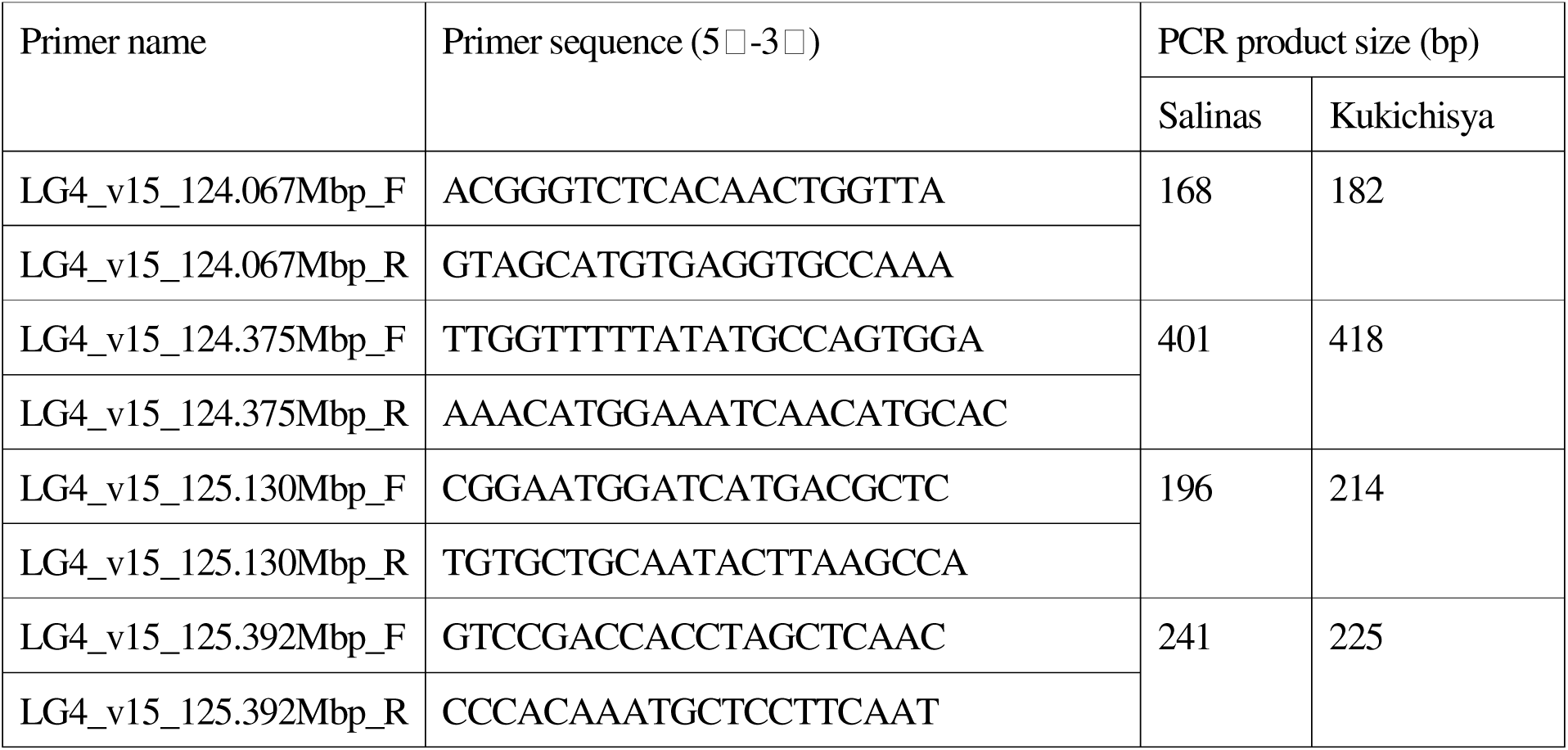
PCR-based markers in the 2-acetyl-1-pyrroline (2AP) locus.

### Whole-genome sequencing analysis of *LsBADH1*

*LsBADH1* gene was sequenced to identify mutations that caused fragrance in ‘Kukichisya.’ The whole-genome sequencing data of ‘Kukichisya’ were aligned against the reference sequence ‘Salinas’ using BWA (Li and Durbin, 2009). The results revealed three synonymous and three non-synonymous mutations in the *LsBADH1* gene (Supplementary Fig. S1); these resulted in amino acid substitution of lysine (K) to arginine (R) at position 17, aspartic acid (D) to glutamic acid (E) at position 65, and serine (S) to glycine (G) at position 295 in *LsBADH1* protein in ‘Kukichisya’ compared with that in ‘Salinas’ (Fig. 3).

**Fig. 3.**
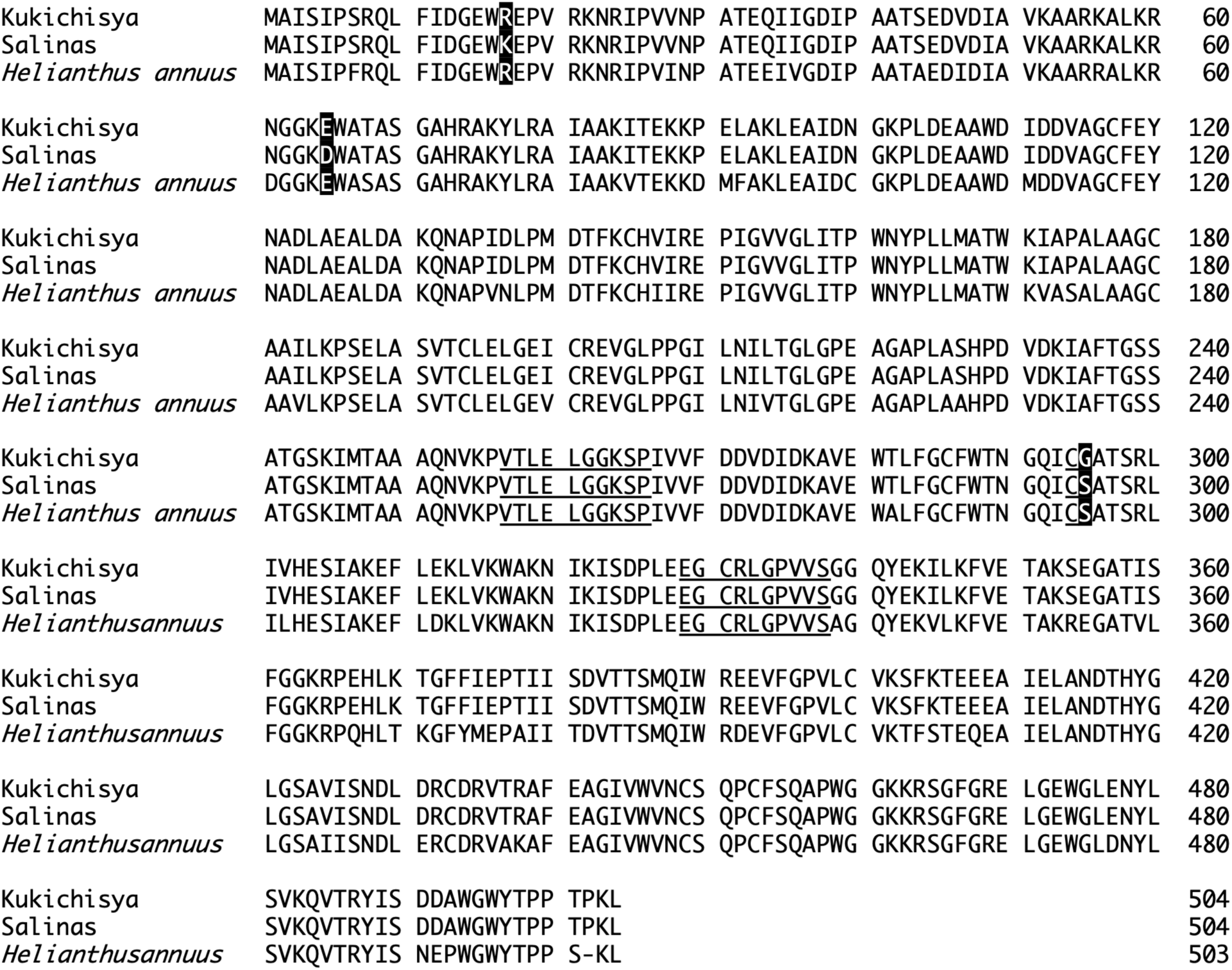
The amino acid sequence of LsBADH1 in fragrant lettuce cultivar: ‘Kukichisya’, non-fragrant lettuce cultivar: ‘Salinas’, and BADH (GenBank No. ACU65243.1) in non-fragrant Sunflower (*Helianthus annuus*). Underlined sequences are believed to be essential for the functional activity of the enzyme. White letters boxed in black indicate non-synonymous single-nucleotide polymorphisms between ‘Kukichisya’ and ‘Salinas’.

The amino acid sequence of the non-synonymous mutations in *LsBADH1* was compared with the BADH protein (LOC110879770) of sunflower (Badouin *et al*., 2017), which belongs to the same Asteraceae family as lettuce. The BADH protein of non-fragrant sunflower and that of the fragrant ‘Kukichisya’ harbor the same amino acid sequences of R at position 17 and E at position 65 (Fig. 3). These results suggested that the amino acid substitution from S to G at position 295 could be mechanism responsible for the functional deficiency of the BADH protein in ‘Kukichisya.’ Additionally, an insertion of 18 bp between exon 5 and 6 was observed in *LsBADH1* (Fig. 4).

**Fig. 4.**
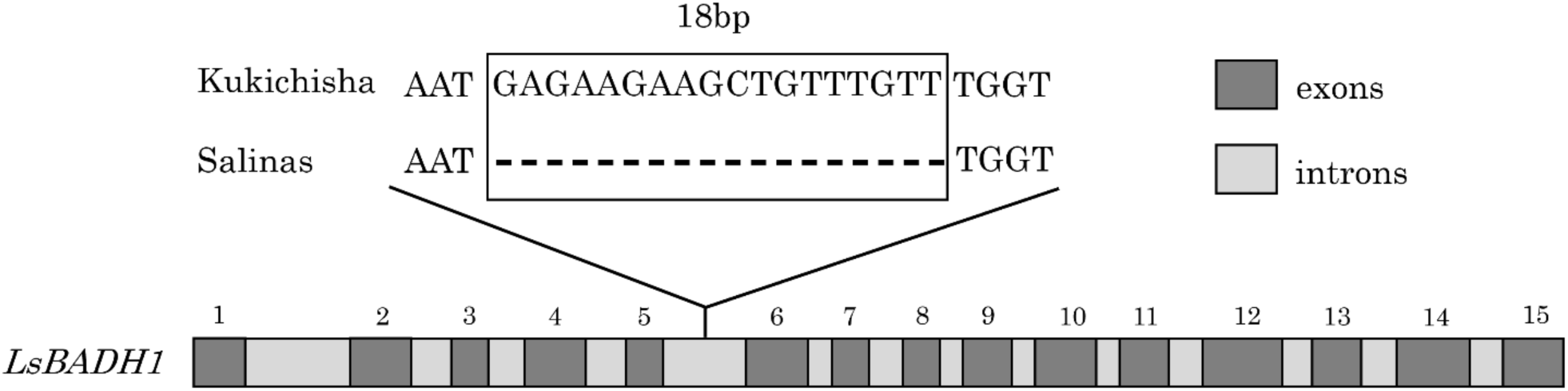
An 18-bp insertion between exon 5 and 6 of *LsBADH1* in ‘Kukichisya’. The insertion is specific to the fragrant cultivar ‘Kukichisya’ and is absent in ‘Salinas’. This polymorphism was utilized to develop a PCR-based marker, providing a robust screening tool for improving the efficiency of fragrant lettuce breeding via marker-assisted selection (MAS).

### Development of PCR-based marker for further mapping of *LsBADH1*

The 18-bp insertion between exons 5 and 6 was used as a nucleotide polymorphism to develop a PCR-based marker to improve the efficiency of fragrant lettuce breeding using marker-assisted selection (MAS) (Fig. 4). The sizes of the PCR products amplified using the *LG4_v15_125.130Mbp* marker were 214 bp for the fragrant cultivar ‘Kukichisya’ and 196 bp for the non-fragrant cultivars (Table 3).

### Base editing of *LsBADH1*

Conventional CRISPR-Cas9 technology typically induces insertions and deletions at the target site, which often result in a frameshift of the coding sequence and expression of truncated or misread peptides. Although this technology can easily disrupt gene function, such abnormal peptides are not favorable in food crops because of the subsequent need to verify their potential to cause allergies. To avoid this, we chose the base editing technology Target-AID, which consists of nuclease-impaired CRISPR-Cas9 and a DNA cytidine deaminase PmCDA1 that enables C to T base substitutions within a range of three bases in the target sequence directed by gRNA (Nishida *et al*, 2016). To install a loss-of-function mutation in the *LsBADH1* gene, gRNA was designed at the 5^th^ exon to generate a premature stop codon at the 170^th^ tryptophan residue (Supplementary Fig. S2). T-DNA vector expressing Target-AID system optimized for dicot plants (Shimatani *et al*, 2017) was introduced via *Agrobacterium* into the lettuce, ‘Oilseed,’ which normally does not produce 2AP. Transformed plants were selected for kanamycin resistance, followed by PCR confirmation. T_0_ and T_1_ individuals were analyzed for editing outcomes, which showed that the only editing pattern was a change from TGG to TAA (Trp170Ter), leading to a truncated protein with 169 amino acids instead of 504 amino acids, as expected. To predict the genotype of the strains (monoallelic or biallelic, heterozygous, or homozygous), eight clones from each individual plant were analyzed by sequencing, showing 87.5% of the edited sequences from T_0_ plants and 100% from one of the T_1_ strains, which was assumed to be biallelic homozygous mutant and used for further analysis. GC-MS analysis demonstrated the presence of 2AP in the edited strain, but not in the wild-type (Fig. 5). These results confirmed that the loss-of-function of *LsBADH* was responsible for the fragrance derived from 2AP.

**Fig. 5.**
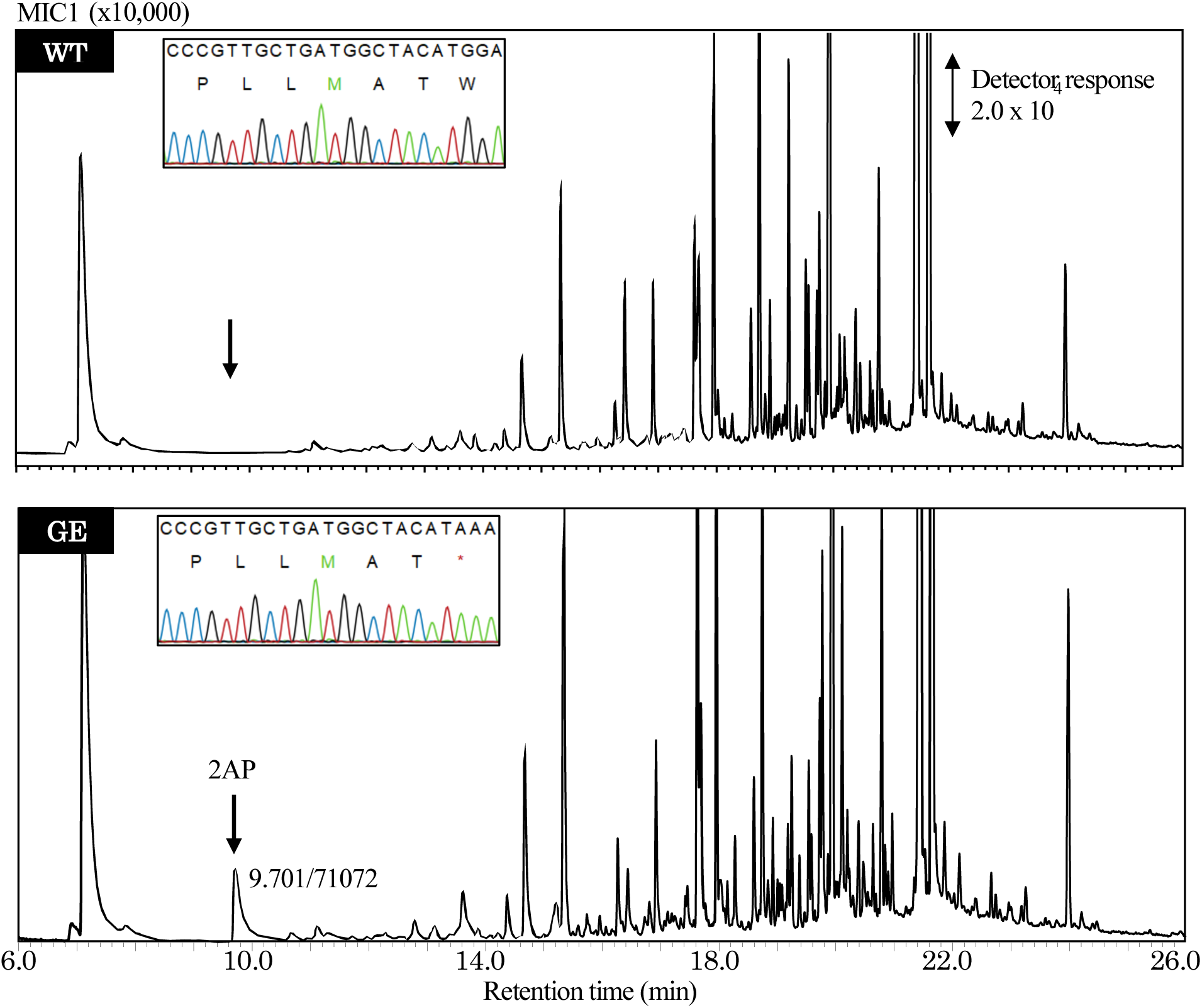
Detection of 2-acetyl-1-pyrroline (2AP) from genome-edited lettuce. An original cultivar ‘Oilseed’ (WT) and genome-edited (GE) LsBADH1(W170*) were used as materials. Extracts from stem and leaf were analyzed by GC-MS. The Sanger sequence electropherograms near top left corner show the target gRNA region. The peak for 2AP is shown using an arrow in the GC-MS chromatogram.

### Confirmation of fragrance-related compounds in ‘Kukichisya’

Validation of volatile compounds of the fragrant cultivar ‘Kukichisya’ and the non-fragrant cultivar ‘Celtuce’ was performed by using GC-MS. Significantly higher levels of the volatile compounds 3-methyl-butanal, 2AP, (*Z*)-(2)-hexen-1-ol, (-)-β-elemene, and octadecanal were detected in ‘Kukichisya’ than in ‘Celtuce.’ 3-methyl-butanal and 2AP were only detected in ‘Kukichisya’ but not in ‘Celtuce’ (Table 1). Moreover, the comparison with the data of the volatile compounds in Pandanus leaves suggested that 2AP was certainly among the fragrance-related compounds in ‘Kukichisya’ and is a key volatile compound that imparted fragrance to ‘Kukichisya’ (Fig. 1).

## Discussion

As a high correlation between 2AP analysis by gas chromatography and organoleptic testing in soybeans has been reported (Juwattanasomran *et al*., 2011), the organoleptic test was adopted for phenotyping of fragrance owing to 2AP in the F_2_ population. Although general fragrances might be thought of as a quantitative trait, 2AP has aspects of qualitative traits because 2AP has a character of a very low odor threshold (Kovach *et al*., 2009). Therefore, the phenotype of the F_2_ population was judged to be fragrant or non-fragrant using the organoleptic test as a qualitative trait in this study. The 1:3 F_2_ segregation ratio of fragrant plants suggested that a single recessive gene was involved (Table 1), and linkage analysis and genetic mapping using PCR-based markers identified the 2AP locus from 124.067 Mbp to 125.392 Mbp in LG4 (Fig. 2). Using whole-genome sequencing, *LOC111877932* encoding BADH located in the 2AP locus was sequenced, resulting in the identification of three nsSNPs (Fig. S1). BADHs are well known for their high aldehyde dehydrogenase activity and wide substrate specificity, and have been reported to exist as two types: BADH1 and BADH2 (Chen *et al*., 2008; Juwattanasomran *et al*., 2011; Liu *et al*., 2015). In soybean, rice, and sorghum, nonfunctional mutations in *BADH1* and *BADH2* are associated with increased 2AP accumulation (Amarawathi *et al*., 2008; Chen *et al*., 2008; Kovach *et al*., 2009; Arikit *et al*., 2011; Juwattanasomran *et al*., 2011; Yundaeng *et al*., 2013; Somta *et al*., 2019). According to the reference genome sequence, lettuce has only a *BADH1* homolog, but no *BADH2* homolog. Although the biochemical pathway of 2AP synthesis has not been fully understood, it is considered that the BADH protein catalyzes the oxidation of a 2AP precursor to produce γ-aminobutyric acid (Chen *et al*., 2008). Because the catalytic residues of Cys-294 has been predicted to interact with the substrate oxygen (Chen *et al*., 2008), we inferred that the non-synonymous mutation Ser295Gly in *LsBADH1* was potentially responsible for the nonfunctional mutation (Fig. 3). Thus, nonfunctional *LsBADH1* with the Ser295Gly mutation mainly caused an accumulation of 2AP in the ‘Kukichisya’ lettuce. Next, Target-AID base editing of genomic *LsBADH1* generated loss-of-function mutants wherein Trp170Ter was successfully installed to delete all catalytic domains and the causative mutation at Ser295, resulting in a complete loss of BADH activity. The homozygous mutant individual showed significant production of 2AP, as detected by GC-MS (Fig 5), demonstrating that mutation in *LsBADH1* was responsible for the accumulation of 2AP.

In the present study, we considered fragrance as a qualitative trait, evaluated it using an organoleptic test (Table 1), and successfully identified the major genes responsible for fragrance. Previous studies have identified minor genes that contribute to the accumulation of 2AP by quantitative trait locus (QTL) analysis of fragrance as a quantitative trait (Amarawathi *et al*., 2008; Somta *et al*., 2019). QTL analysis using quantitative data on fragrance traits may allow the identification of additional genes that contribute to the accumulation of 2AP in lettuce. Furthermore, exogenous application of some elements, such as manganese (Mn), zinc (Zn), and silicon (Si), has been reported to be effective in increasing 2AP formation in fragrant rice (Li *et al*., 2016; Mo *et al*., 2017; Luo *et al*., 2019). Proline, which is a precursor of 2AP, has also been demonstrated to increase 2AP content upon foliar spraying the initial heading stage of fragrant rice (Luo *et al*., 2020). The biologically meaningful prediction that the accumulation of 2AP in lettuce could be increased by different cultivation approaches is worth exploring as a production system for fragrant lettuce cultivars.

The primary sources of fragrances are volatile compounds with molecular weights below 300 Da and high vapor pressures (Kumar et al. 2018). Because fragrant cultivars can increase consumer acceptance and command higher prices worldwide, they are of interest to the food industry as high-quality crops. Notably, fragrance due to 2AP is also associated with important flavors in a variety of cooked foods, such as popcorn, corn tortillas, and bread crust (Adams and De Kimpe, 2006). 2AP accumulates in the aerial plant organs (e.g., leaves, stems, and seeds) (Kovach *et al*., 2009), and is released by cooking methods such as boiling or baking (Buttery *et al*., 1983). For rice and soybeans, both fragrant and non-fragrant cultivars are cooked using the same methods. In the case of lettuce, the fragrant cultivars are typically eaten not only as fresh salads but also after cooking by heat treatment for the exploitation of fragrance (Starkenmann *et al*., 2019). As the present lettuce cultivars were selected for fresh salads without heat treatment, many cultivars may not match expectations for cooking with heat treatment. It is necessary to breed new cultivars that are suitable for cooking, rather than backcrossing the present cultivars using a selection marker (Seki *et al*., 2024).

This study demonstrated that the loss of *LsBADH1* function results in the sweet fragrance phenotype in lettuce through the accumulation of 2AP. However, the significance of this gene goes far beyond simply adding flavor. Introducing fragrance into lettuce has the potential to fundamentally reshape breeding strategies that have long been optimized exclusively for fresh salad consumption. Disruption of *LsBADH1* not only creates a novel sensory trait, but also has the potential to redefine lettuce as a crop with value in cooked and processed culinary applications. In this context, a single gene has the capacity to alter flavor, thereby transforming the breeding objectives and culinary identity of lettuce. Therefore, our findings represent more than the identification of a fragrance-associated gene; they provide a foundation for generating new diversity and creating previously unexplored value in lettuce as a vegetable crop.

## Supplementary Data

The following supplementary data are available at JXB online.

Fig. S1. The coding sequence of *LsBADH1* in non-fragrant lettuce cultivar ‘Salinas’ and fragrant lettuce cultivar ‘Kukichisya’.

Fig. S2. *LsBADH* nucleotide sequence and gRNA design for base editing.

## Acknowledgments

The authors would like to thank Dr. Ushio Fujikura and Dr. Kanami Yamaguchi for their assistance with the genome editing experiments. The authors thank Satoshi Kawamoto and Nanako Fukushima for their technical assistance with the Sanger sequencing.

## Author contributions

KS and YU conducted experiments. KS developed mapping populations. KS performed linkage, fine mapping, phylogenetic, and resequencing analyses. MY, KN, RK, and YU performed the genome editing. KM performed GC-MS analysis of the fragrance-related compounds. KS, KM, and YU wrote the manuscript. All the authors have read and approved the final manuscript.

## Data availability

All data generated or analyzed during this study are included in this published article.

