## Supplemental Figures for "*LsBADH1* is responsible for sweet fragrance in lettuce (*Lactuca sativa* L.) through 2-acetyl-1-pyrroline biosynthesis"

### Supplementary Figures

```
                                                               *
Kukichisya    ATGGCGATCTCGATACCATCGCGTCAGTTATTCATAGACGGAGAATGGAGGGAGCC    56
Salinas       ATGGCGATCTCGATACCATCGCGTCAGTTATTCATAGACGGAGAATGGAAGGAGCC    56

Kukichisya    TGTTAGGAAGAATCGAATCCCTGTCGTAAATCCCGCCACCGAGCAGATCATCGGGG    112
Salinas       TGTTAGGAAGAATCGAATCCCTGTCGTAAATCCCGCCACCGAGCAGATCATCGGGG    112

Kukichisya    ATATCCCAGCAGCTACATCTGAAGATGTTGATATTGCTGTGAAAGCAGCTCGTAAA    168
Salinas       ATATCCCAGCAGCTACATCTGAAGATGTTGATATTGCTGTGAAAGCAGCTCGTAAA    168
                                        *     *
Kukichisya    GCTCTTAAACGCAATGGAGGAAAAGAGTGGGCTACAGCTTCTGGAGCACATCGTG    223
Salinas       GCTCTTAAACGCAATGGAGGAAAAGACTGGGCAACAGCTTCTGGAGCACATCGTG    223

Kukichisya    CCAAGTATCTGCGTGCCATTGCTGCAAAGATAACTGAGAAAAAACCGGAATTAGCA    279
Salinas       CCAAGTATCTGCGTGCCATTGCTGCAAAGATAACTGAGAAAAAACCGGAATTAGCA    279
                                     *
Kukichisya    AAACTTGAAGCCATTGATAATGGAAAACCACTTGATGAAGCAGCATGGGATATAGA    335
Salinas       AAACTTGAAGCCATTGATAATGGCAAACCACTTGATGAAGCAGCATGGGATATAGA    335
                                            *
Kukichisya    TGATGTGGCTGGATGTTTTGAATATAATGCTGATCTTGCTGAAGCTTTGGATGCTAA    392
Salinas       TGATGTGGCTGGATGTTTTGAATATAATGCCGATCTTGCTGAAGCTTTGGATGCTAA    392

Kukichisya    GCAAAACGCACCTATTGATCTTCCAATGGATACATTCAAATGTCATGTGATTAGGGA    449
Salinas       GCAAAACGCACCTATTGATCTTCCAATGGATACATTCAAATGTCATGTGATTAGGGA    449

Kukichisya    ACCTATTGGTGTTGTTGGGCTGATTACTCCATGGAATTACCCGTTGCTGATGGCTAC    506
Salinas       ACCTATTGGTGTTGTTGGGCTGATTACTCCATGGAATTACCCGTTGCTGATGGCTAC    506

Kukichisya    ATGGAAAATTGCACCTGCCCTGGCTGCTGGTTGTGCTGCAATACTTAAGCCATCAG    562
Salinas       ATGGAAAATTGCACCTGCCCTGGCTGCTGGTTGTGCTGCAATACTTAAGCCATCAG    562

Kukichisya    AATTGGCATCTGTGACATGCCTGGAATTAGGTGAAATATGCAGGGAGGTAGGTCTT    618
Salinas       AATTGGCATCTGTGACATGCCTGGAATTAGGTGAAATATGCAGGGAGGTAGGTCTT    618

Kukichisya    CCCCCTGGTATTCTCAATATCCTAACTGGATTAGGTCCAGAAGCAGGTGCCCCTTT    674
Salinas       CCCCCTGGTATTCTCAATATCCTAACTGGATTAGGTCCAGAAGCAGGTGCCCCTTT    674

Kukichisya    GGCTTCTCACCCTGATGTTGATAAGATTGCATTTACAGGAAGCAGTGCCACAGGAA    730
Salinas       GGCTTCTCACCCTGATGTTGATAAGATTGCATTTACAGGAAGCAGTGCCACAGGAA    730

Kukichisya    GCAAGATTATGACTGCAGCAGCTCAAAATGTCAAGCCTGTTACACTTGAGCTTGGT    786
Salinas       GCAAGATTATGACTGCAGCAGCTCAAAATGTCAAGCCTGTTACACTTGAGCTTGGT    786

Kukichisya    GGAAAGAGTCCAATAGTTGTGTTTGATGATGTTGACATTGATAAAGCTGTTGAGTG    842
Salinas       GGAAAGAGTCCAATAGTTGTGTTTGATGATGTTGACATTGATAAAGCTGTTGAGTG    842
                                                      *
Kukichisya    GACACTCTTTGGTTGCTTCTGGACAAATGGGCAAATCTGCGGTGCAACTTCTCGAC    898
Salinas       GACACTCTTTGGTTGCTTCTGGACAAATGGGCAAATCTGCAGTGCAACTTCTCGAC    898

Kukichisya    TCATAGTGCATGAAAGTATTGCTAAGGAGTTTTTAGAGAAGCTTGTGAAGTGGGCT    954
Salinas       TCATAGTGCATGAAAGTATTGCTAAGGAGTTTTTAGAGAAGCTTGTGAAGTGGGCT    954

Kukichisya    AAAAATATCAAGATTTCAGATCCATTAGAAGAAGGTTGCAGGCTTGGACCTGTTGT    1010
Salinas       AAAAATATCAAGATTTCAGATCCATTAGAAGAAGGTTGCAGGCTTGGACCTGTTGT    1010

Kukichisya    TAGTGGTGGACAGTATGAGAAGATATTGAAGTTTGTTGAAACTGCCAAAAGCGAAG    1066
Salinas       TAGTGGTGGACAGTATGAGAAGATATTGAAGTTTGTTGAAACTGCCAAAAGCGAAG    1066

Kukichisya    GTGCAACCATTTCATTTGGAGGAAAACGTCCCGAGCATTTGAAAACCGGATTTTTT    1122
Salinas       GTGCAACCATTTCATTTGGAGGAAAACGTCCCGAGCATTTGAAAACCGGATTTTTT    1122

Kukichisya    ATCGAGCCAACAATCATTAGTGATGTCACCACATCCATGCAAATTTGGAGAGAGGA    1178
Salinas       ATCGAGCCAACAATCATTAGTGATGTCACCACATCCATGCAAATTTGGAGAGAGGA    1178

Kukichisya    AGTTTTTGGACCTGTTTTATGCGTAAAATCATTTAAAACAGAAGAAGAAGCAATCGA    1235
Salinas       AGTTTTTGGACCTGTTTTATGCGTAAAATCATTTAAAACAGAAGAAGAAGCAATCGA    1235

Kukichisya    ATTAGCAAATGACACCCATTATGGTCTGGGTTCTGCTGTTATATCCAACGATTTGGA    1292
Salinas       ATTAGCAAATGACACCCATTATGGTCTGGGTTCTGCTGTTATATCCAACGATTTGGA    1292

Kukichisya    CCGGTGTGATCGTGTGACAAGGGCTTTTGAGGCAGGTATTGTTTGGGTAAACTGCT    1348
Salinas       CCGGTGTGATCGTGTGACAAGGGCTTTTGAGGCAGGTATTGTTTGGGTAAACTGCT    1348

Kukichisya    CCCAGCCTTGCTTCTCTCAAGCTCCGTGGGGTGGGAAAAAGCGTAGTGGATTTGG    1403
Salinas       CCCAGCCTTGCTTCTCTCAAGCTCCGTGGGGTGGGAAAAAGCGTAGTGGATTTGG    1403

Kukichisya    TCGTGAACTAGGAGAATGGGGACTTGAGAATTATTTGAGCGTAAAGCAGGTGACTC    1459
Salinas       TCGTGAACTAGGAGAATGGGGACTTGAGAATTATTTGAGCGTAAAGCAGGTGACTC    1459

Kukichisya    GTTATATTTCTGATGATGCTTGGGGTTGGTATACGCCTCCAACTCCTAAGCTTTGA    1515
Salinas       GTTATATTTCTGATGATGCTTGGGGTTGGTATACGCCTCCAACTCCTAAGCTTTGA    1515
```

Fig. S1. The coding sequence of *LsBADH1* in non-fragrant lettuce cultivar: ‘Salinas’, and fragrant lettuce cultivar: ‘Kukichisya’. ‘Kukichisya’ possesses six single nucleotide polymorphisms (SNPs) as indicated by the asterisks.

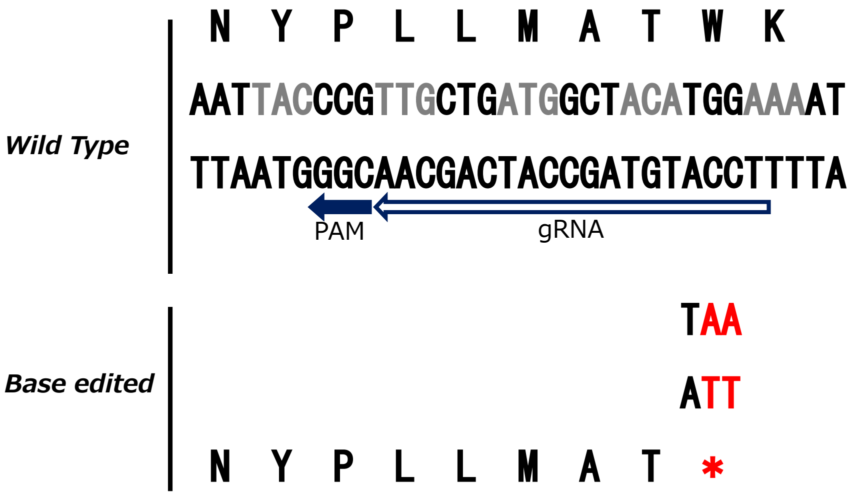


Fig. S2. *LsBADH* nucleotide sequence and gRNA design for base editing. The 32 bp of sense and antisense sequences of the fifth exon of *LsBADH1* is shown. The black and white arrows, respectively, indicate the PAM and gRNA sequences in the antisense strand, which is intended to install a stop codon at W170, as the Target-AID base editing induces a C to T base change efficiently at 16 to 18 base position upstream of the PAM sequence.
